# Long-term culturing of porcine nodose ganglia

**DOI:** 10.1101/706945

**Authors:** Shin-Ping Kuan, Kalina R. Atanasova, Maria V. Guevara, Emily N. Collins, Leah R. Reznikov

## Abstract

**Background:** Neuronal cell cultures are widely used in the field of neuroscience. Cell dissociation allows for the isolation of a desired cell type, yet the complexity that distinguishes the nervous system is often lost as a result. Thus, culturing neural tissues in *ex vivo* format provides a physiological context that more closely resembles the *in vivo* environment. Nodose ganglia neurons have been extensively studied both in dissociated form and acutely in slice format. However, methods to culture long-term *ex vivo* have not been established.

**New Method:** We developed a simple method to culture nodose ganglia neurons from neonatal pigs long-term in *ex vivo* format using an in-house media formulation derived from commercially available components.

**Results:** Cultures were viable for approximately 12 months. mRNA expression for nestin, a marker of neural progenitor cells, was stable across time. Vasoactive intestinal peptide and tachykinin, markers of nodose neurons, showed either no statistically significant differences or decreased across time, respectively. mRNA expression for glia fibrillary acidic protein and myelin basic protein showed no statistically significant differences over time.

**Comparison with Existing Method(s):** There are currently no methods that describe long-term culturing of porcine nodose ganglia. Further, the media formulation we developed is new and not previously reported.

**Conclusions:** The simple procedure we developed for culturing nodose ganglia will enable both short-term and long-term investigations aimed at understanding peripheral ganglia *in vitro*. It is also possible that the methods described herein can be applied to other animal models, adult samples, and other neural tissues.

## 1. Introduction

Primary neuronal cultures are powerful model systems to dissect nervous system function, development, and cellular processes [1]. They can closely mimic *in vivo* characteristics of the parent tissue in an isolated environment. [2]. However, there are several challenges encountered during preparation of primary cultures. These include, but are not limited to, the following: 1) number of cells available; 2) separation of cells of interests from other cells present in originating tissue; 3) viability of cells post isolation; and 4) loss of *in vivo* cytoarchitecture and cellular complexity.

Although numerous techniques have been developed to culture primary neurons [1], one approach that circumvents some of these challenges is the organotypic slice culture [2]. Slice cultures preserve many of the structural and synaptic elements of the original tissue, resulting in a more *in vivo* like state. Still, even in organotypic slice cultures, neuron survival is a challenge. Survival of neurons is dependent upon age of the subject, media composition, and health of the donor [2]. Media composition has advanced with the introduction of neurobasal media and B-27 supplement, the combination of which prolong neuronal survival *in vitro* [3]. Yet even with media refinement, neuron survivability on average ranges from weeks [4] to months [5].

Many of the aforementioned methods have been developed and/or refined for rodent central nervous system neurons. However, there is continued interest in the peripheral nervous system, as well as appeal in utilizing non-rodent models [6]. Thus, methods that enable the culturing of peripheral ganglia from non-rodent model systems could fill a critical gap in our knowledge. Further, refining methods to enable long-term culturing of neurons would greatly facilitate studies aimed at delineating neuronal plasticity, age-related processes, and therapeutics.

## 2. Materials and Methods

### 2.1 Animals

Newborn pigs (~1 week of age, n=4, male, n = 5 female) were used for this study. Pigs were sedated and anesthetized with ketamine and xylazine. Under sedation, intravenous Euthasol (Virbac) was delivered for euthanasia. Ganglia (right and left) from each pig were collected following protocols previously described [6]. The University of Florida Animal Care and Use Committee approved all procedures.

### 2.2 Culturing

Twelve of the collected ganglia were placed on a PET track-etched membrane with a 1.0 μm pore (Falcon, Cat. #353102) in a 6-well plate and placed in a water-jacketed incubator with 5% CO2. Cultures were maintained at 37 °C. This method is similar to that described by Stoppini [7]. Media was changed every 2-3 days (e.g., on Mondays and Thursdays) with new 1.5-2.0 mls of media added basolateral to the membrane. Six of the ganglia were immediately fixed in 4% paraformaldehyde (n = 3 animals) or placed in RNAlater (ThermoFisher, Cat. #AM7021) for RNA extraction.

### 2.3 Media

An in-house media consisting of the following components was developed: 49 mls of Neurobasal A - Glutamine (GIBCO, ThermoFisher Cat. #10888022); 49 mls of Ham’s F-12 Nutrient Mix + Glutamine (GIBCO, ThermoFisher Cat. #11765054); 1 ml of B-27 Supplement (GIBCO, ThermoFisher Cat. #17504044); 1 ml of Penicillin-Streptomycin solution (GIBCO, ThermoFisher, Cat. #15140122); 0.5 mls of CultureOne Supplement (GIBCO, ThermoFisher Cat. #A3320201). Media was sterile filtered with a 0.2 μm filter (Corning, ThermoFisher, Cat. #430049) and stored at 4 °C.

### 2.4 RNA isolation and qRT-PCR

RNA from the whole nodose ganglia was isolated using RNeasy Lipid Tissue kit (Qiagen, Cat. #74804) with optional DNase digestion. RNA concentrations were assessed using a NanoDrop spectrophotometer. RNA was reverse transcribed (100 ng) using Superscript VILO Master Mix (ThermoFisher, Cat. #11755050). Briefly, RNA and master mix were incubated for 10 mins at 25 °C, followed by 60 mins at 42°C, and 5 mins at 85°C. Transcript abundance for nestin, vasoactive intestinal peptide (VIP), glia fibrillary acidic protein (GFAP), myelin basic protein (MBP), and tachykinin (TAC) were measured using methods similar to those previously described [8, 9]. Actin was used as a housekeeping gene. Primer sequences are shown in Table 1. All qRT-PCR data were acquired using fast SYBR green master mix (Applied Biosystems, Cat. #4385618) and a LightCycler 96 (Roche). Standard ΔΔCT methods were used for analysis.

### 2.5 Immunofluorescence

Cultured and non-cultured ganglia were placed in 2% paraformaldehyde solution (Electron Microscopy Sciences, 8% stock, Cat. #1578100) for 4 hours and then sucrose-protected at 4 °C in 30% sucrose for 72 hours. Tissues were sectioned at a thickness of 10μm and mounted onto SuperFrost Plus microscope slides (ThermoFisher Scientific, Cat. #22-037-246).

Immunofluorescence procedures were similar to those previously described [8]. Briefly, cross-sections from a single cohort of pigs were selected and fixed in 2% paraformaldehyde for 15 minutes. Tissues were then permeabilized in 0.15% Triton X-100, followed by blocking in PBS Superblock (ThermoFisher Scientific, Cat. #37515) containing 4% normal goat serum (Jackson Laboratories, Cat #005-000-121). Tissues were incubated with rabbit anti-neurofilament heavy chain antibody (Novus Biologicals, Cat. #NB300-135) at 1:1000 dilution for 2 hours at 37 °C. Tissues were washed thoroughly in PBS and incubated with Alexa fluor 568-conjugated goat anti-rabbit secondary antibody (ThermoFisher, Cat. #A11011) at a 1:1000 dilution for 1 hour at room temperature. Tissues were washed and a Hoechst stain was performed as previously described [8]. A 1:10 glycerol/PBS solution was used to cover the sections and cover glass added. Sections were imaged on a Zeiss Axio Zoom V16 microscope.

### 2.6 Statistical Analysis

A one-way ANOVA was used to assess the effect of time for a given gene, followed by a Tukey multiple comparison test. All tests were carried out using GraphPad Prism 7.0a. Statistical significance was determined as P < 0.05.

## 3. Results

Ganglia were placed on semi-permeable inserts (Figure 1A) and cultured for 6 or 12 months. We first examined mRNA transcript abundances for markers neural progenitors, mature neurons, and glial cells. Nestin, a marker of neural progenitors [10], was relatively stable at the time points measured (Figure 1B). Both the astrocytic marker GFAP and the Schwann cell marker MBP [9] showed non-significant decreases across time (Figure 1B). We also examined markers of porcine nodose ganglia neurons [11], VIP and tachykinin, a precursor for substance P. Although VIP appeared to increase across time, this elevation was not statistically significant (Figure 1B). In contrast, expression of tachykinin decreased over time (Figure 1B). We also confirmed presence of neurons through by examining neurofilament protein expression in nodose ganglia cross sections [12] (Figure 1C, 1D). Neurofilament-positive fibers and neuronal cell bodies were observed at 12 months, although there were qualitatively fewer cell nuclei and neurofilament-positive cell bodies.

**Fig. 1.**
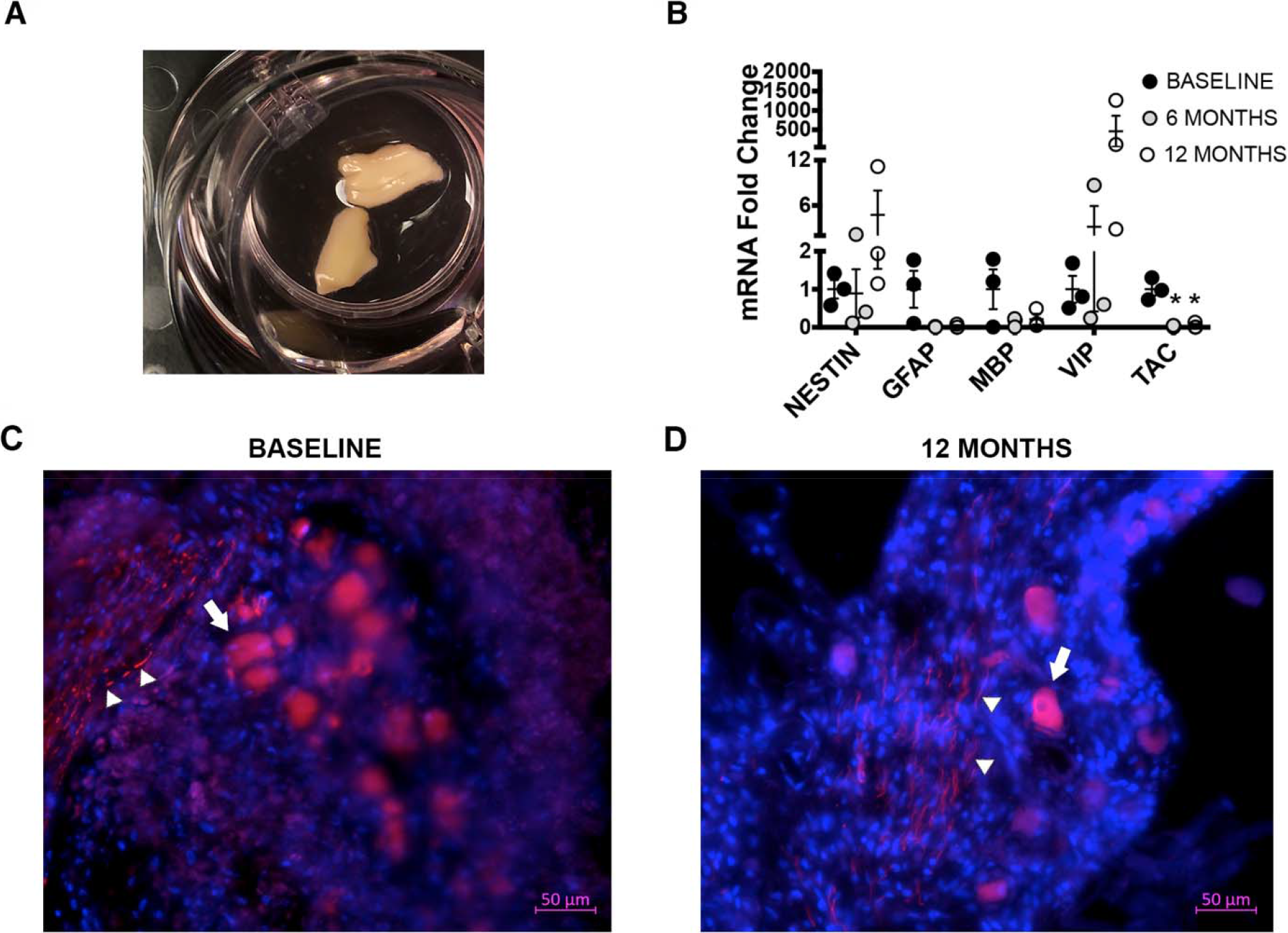
Properties of nodose ganglia neurons cultured long-term. A) Ganglia on semi-permeable inserts. B) Transcript abundance for nestin, glia fibrillary acidic protein (GFAP), myelin basic protein (MBP), vasoactive intestinal peptide (VIP) and tachykinin (TAC). All levels are expressed as fold change relative to baseline values. Mean ± S.E.M shown. Neurofilament labeling (in red) in nodose ganglia at baseline (C) and after 12 months (D) culturing. Arrow heads highlight examples of nerve fibers and arrows represent examples of cell bodies. Nuclei are shown in blue. * P < 0.05 compared to baseline levels.

## 4. Discussion

In the current study, we demonstrated simple methods for long-term culture of porcine nodose ganglia. We observed preservation of neuronal markers both at the mRNA and protein levels after 12 months of culturing, representing a significant advancement for the field. We anticipate these methods will be of broad interest because pigs are increasingly used as models of human disease, and because the nodose ganglia house sensory neurons of multiple visceral organs [6].

The media formulation we designed included numerous compounds. We are not certain which component(s) were critical in maintaining neurons long-term *in vitro*. A side-by-side comparison of neurobasal A media with Ham’s F-12 nutrient mix suggest numerous possible players. For example, we suspect that biotin, lipoic acid, linoleic acid and putrescine were involved, as these are found in B-27 supplement [13]. We also anticipate that the CultureOne supplement facilitated long-term maintenance of neurons.

Our study offers several advantages. For example, magnetic resonance imaging suggest that the neuronal maturation and dendritic arborization of a one week old piglet brain is equivalent to a one month old human brain [14]. This suggests that the method described here might be of value and applicable to human tissues, and possibly adult animals. Further, the use of the semi-permeable insert gives rise to the possibility of co-culture experiments, with nodose ganglia in the top compartment and tissue of interest in the bottom compartment. A final advantage is that the procedure we developed to culture neurons long-term is simple and relatively low maintenance.

Our study also has limitations. We are not certain whether the media formulation we developed is superior to other formulations. Additionally, as mentioned, we do not know which specific component(s) in the media facilitates long-term survival of neurons. Such studies might be costly and/or time-consuming to perform, although potentially of high impact. We also do not know whether the phenotypic cell complexity observed *in vivo* is maintained long-term *in vitro*. The decrease in tachkyinin expression might suggest not.

In summary, we developed simple procedures for long-term culturing of porcine nodose ganglia. It is our hope that these methods will have broad range applicability to additional species, other peripheral and central nervous system tissues, and different developmental time points.

## Acknowledgements

We thank Jonai Moore, Kevin Vogt, Yan-Shin Liao and Joshua Dadural for helpful comments and technical assistance. This work was supported by the National Institutes of Health R00HL119560 (PI, LRR) and 10T2TR001983 (Co-I, LRR).

